# Susceptibility of sheep to experimental co-infection with the ancestral lineage of SARS-CoV-2 and its alpha variant

**DOI:** 10.1101/2021.11.15.468720

**Authors:** Natasha N. Gaudreault, Konner Cool, Jessie D. Trujillo, Igor Morozov, David A. Meekins, Chester McDowell, Dashzeveg Bold, Mariano Carossino, Velmurugan Balaraman, Dana Mitzel, Taeyong Kwon, Daniel W. Madden, Bianca Libanori Artiaga, Roman M. Pogranichniy, Gleyder Roman-Sosa, William C. Wilson, Udeni B. R. Balasuriya, Adolfo García-Sastre, Juergen A. Richt

## Abstract

Severe acute respiratory syndrome coronavirus 2 (SARS-CoV-2) is responsible for a global pandemic that has had significant impacts on human health and economies worldwide. SARS-CoV-2 is highly transmissible and the cause of coronavirus disease 2019 (COVID-19) in humans. A wide range of animal species have also been shown to be susceptible to SARS-CoV-2 infection by experimental and/or natural infections. Domestic and large cats, mink, ferrets, hamsters, deer mice, white-tailed deer, and non-human primates have been shown to be highly susceptible, whereas other species such as mice, dogs, pigs, and cattle appear to be refractory to infection or have very limited susceptibility. Sheep (*Ovis aries*) are a commonly farmed domestic ruminant that have not previously been thoroughly investigated for their susceptibility to SARS-CoV-2. Therefore, we performed *in vitro* and *in vivo* studies which consisted of infection of ruminant-derived cell cultures and experimental challenge of sheep to investigate their susceptibility to SARS-CoV-2. Our results showed that sheep-derived cell cultures support SARS-CoV-2 replication. Furthermore, experimental challenge of sheep demonstrated limited infection with viral RNA shed in nasal and oral swabs primarily at 1-day post challenge (DPC), and also detected in the respiratory tract and lymphoid tissues at 4 and 8 DPC. Sero-reactivity was also observed in some of the principal infected sheep but not the contact sentinels, indicating that transmission to co-mingled naïve sheep was not highly efficient; however, viral RNA was detected in some of the respiratory tract tissues of sentinel animals at 21 DPC. Furthermore, we used challenge inoculum consisting of a mixture of two SARS-CoV-2 isolates, representatives of the ancestral lineage A and the B.1.1.7-like alpha variant of concern (VOC), to study competition of the two virus strains. Our results indicate that sheep show low susceptibility to SARS-CoV-2 infection, and that the alpha VOC outcompeted the ancestral lineage A strain.

## Introduction

Severe acute respiratory syndrome coronavirus 2 (SARS-CoV-2) continues to significantly impact human health and the economies of countries around the world. Since its identification in December 2019, the virus has rapidly spread and evolved resulting in the emergence of multiple variants. Several variants of concern (VOCs) have been shown to be more infectious/transmissible in humans and at the same time reduce the efficacy of currently available vaccines (1–3). Furthermore, SARS-CoV-2 is a zoonotic virus with a wide-range of susceptible animal species which could serve as secondary reservoirs to perpetuate viral evolution and produce novel variants (4–6). As of 31 October 2021, the OIE reports that more than 598 natural infections have been identified in 14 different animal species including companion animals such as: cats and dogs; zoo animals including large cats, otter and gorillas; and farmed or wild animals, including mink and white-tailed deer (www.oie.int); (7). Experimental infections have demonstrated non-human primates, hamsters, ferrets, cats, and deer to be readily susceptible species, while dogs, pigs and cattle appear to have limited susceptibility, and avian species such as chickens and ducks are resistant to infection (8–17). Rabbits, raccoon dogs, fruit bats, several wild mice species, and skunks have also been shown to be susceptible after experimental infection with SARS-CoV-2 (10, 18–20). Mice are not susceptible to the ancestral lineage A strains but to the alpha, beta and gamma VOCs with the 501Y mutation in the Spike protein (21, 22).

Transmission of SARS-CoV-2 occurs via respiratory droplets/aerosols and direct contact with infected individuals or indirect via contaminated fomites. Host factors which contribute to transmission include duration of infection, intensity of shedding, and host behavior and population density. Interactions regularly occur between infected humans and animals, infected animals with other animals, and their environments, complicating the epidemiology of SARS-CoV-2. Natural SARS-CoV-2 infection in farmed mink has demonstrated the significant economic and public health repercussions that can result when variants emerge from an animal reservoir (4, 23). Cross-species maintenance of SARS-CoV-2 makes control of the virus exponentially more complicated. Identification of susceptible hosts and respective biosurveillance is critical to mitigate future secondary zoonotic events. Furthermore, the absence or limited manifestation of clinical symptoms presented by some of the highly susceptible species, such as cats, ferrets or wild-tailed deer, likely contributes to many unnoticed and underreported zoonotic and reverse-zoonotic events.

Sheep are economically important domestic ruminants, commonly farmed worldwide. Sheep are often maintained in large flocks, and frequently interact with humans during standard farming activities, sheering, milking, slaughter for meat, at petting-zoos; sheep also potentially have contact with other susceptible animal species such as mice, cats and deer. However, to date there has been very limited data on the susceptibility of sheep to SARS-CoV-2. An *in silico* study modeling the interactions between species-specific ACE2 receptors and the SARS-CoV-2 spike protein (24) indicates that sheep are potentially susceptible to SARS-CoV-2. In agreement with that study, replication of SARS-CoV-2 in sheep-derived respiratory tissues was demonstrated in an *ex vivo* infection study (25). However, a limited serological survey of sheep in close contact with humans pre- and post-pandemic at a veterinary campus in Spain discovered no detectable SARS-CoV-2 antibodies, suggesting sheep might not be readily susceptible (26).

In this study, we investigated the susceptibility of sheep (*Ovis aries*) both *in vitro* and *in vivo* to SARS-CoV-2. First, ruminant-derived cell cultures were infected and viral growth kinetics determined. Secondly, we challenged eight sheep and studied virus transmission by co-mingling two naïve sentinel sheep at 1-day post challenge (DPC). *Postmortem* evaluations and tissues collections were performed at 4, 8 and 21 DPC to determine SARS-CoV-2 infection and disease progression. Clinical (swabs) and serological samples were collected over the course of the study to monitor viral shedding and seroconversion, respectively. In addition, sheep were inoculated with a mixture of two SARS-CoV-2 strains, representatives of the ancestral lineage A strain and the a B.1.1.7-like alpha variant of concern (VOC) to study the competition of the two viruses. The results of this study are important for understanding the role of sheep in the ecology of SARS-CoV-2.

## Materials and methods

### Cells and virus isolation/titrations

Vero E6 cells (ATCC; Manassas, VA) and Vero E6 cells stably expressing transmembrane serine protease 2 (Vero-E6/TMPRSS2) (27) were obtained from Creative Biogene (Shirley, NY) via Kyeong-Ok Chang at KSU and used for virus propagation and titration. Cells were cultured in Dulbecco’s Modified Eagle’s Medium (DMEM, Corning, New York, N.Y, USA), supplemented with 10% fetal bovine serum (FBS, R&D Systems, Minneapolis, MN, USA) and antibiotics/antimycotics (ThermoFisher Scientific, Waltham, MA, USA), and maintained at 37 °C under a 5% CO_2_ atmosphere. The addition of the selection antibiotic, G418, to cell culture medium was used to maintain TMPRSS2 expression but was not used during virus cultivation or assays. The SARS-CoV-2/human/USA/WA1/2020 lineage A (referred to as lineage A WA1; BEI item #: NR-52281) and SARS-CoV-2/human/USA/CA_CDC_5574/2020 lineage B.1.1.7 (alpha VOC B.1.1.7; NR-54011) strains were acquired from BEI Resources (Manassas, VA, USA). A passage 2 plaque-purified stock of lineage A WA1 and a passage 1 of the alpha VOC B.1.1.7 stock were used for this study. Virus stocks were sequenced by next generation sequencing (NGS) using the Illumina MiSeq and the consensus sequences were found to be homologous to the original strains obtained from BEI (GISAID accession numbers: EPI_ISL_404895 (WA-CDC-WA1/2020) and EPI_ISL_751801 (CA_CDC_5574/2020).

To determine infectious virus titers of virus stocks and study samples, 10-fold serial dilutions were performed on Vero-E6/TMPRSS2 cells. The presence of cytopathic effects (CPE) after 96 hours incubation at 37 °C was used to calculate the 50% tissue culture infective dose (TCID_50_)/ml using the Spearman-Kaerber method (28). Virus isolation attempts were only performed on samples with ≥10^3^ RNA copy number per mL, as this was our approximate limit of detection (LOD) for viable virus using this method. Virus isolation was performed by culturing 100 µl of filtered (0.2 µm; MidSci, St. Louis, MO) sample /well in duplicate on Vero E6/TMPRRS2 cells and monitoring for CPE for up to 5 days post inoculation.

### Susceptibility of ovine and bovine cells to SARS-CoV-2

The SARS-CoV-2 USA-WA1/2020 strain was passaged 3 times in Vero-E6 cells to establish a stock virus for infection experiments. Primary ovine kidney and American pronghorn lung cells (provided by USDA ARS-ABADRU), and bovine fetal fibroblast, Madin-Darby ovine kidney and Madin-Darby bovine cell lines (ATCC; Manassas, VA) were infected at approximately 0.1 multiplicity of infection (MOI). Infected cell supernatants were collected at 0, 2, 4, 6 or 8 days post infection (DPI) and stored at −80°C until further analysis. Cell lines were tested in at least two independent infection experiments. Cell supernatants were titrated on Vero E6 cells to determine SARS-CoV-2 replication kinetics by virus titers (TCID_50_/mL).

### Ethics statement

All animal studies and experiments were approved and performed under the Kansas State University (KSU) Institutional Biosafety Committee (IBC, Protocol #1460) and the Institutional Animal Care and Use Committee (IACUC, Protocol #4508.2) in compliance with the Animal Welfare Act. All animal and laboratory work were performed in biosafety level-3+ and −3Ag laboratories and facilities in the Biosecurity Research Institute at KSU in Manhattan, KS, USA.

### Virus challenge of animals

Ten male sheep, approximately 6 months of age, were acquired from Frisco Farms (Ewing, IL) and acclimated for ten days in BSL-3Ag biocontainment with feed and water *ad libitum* prior to experimental procedures. On day of challenge, eight principal infected sheep were inoculated with a 1:10 titer ratio of lineage A WA1 and the alpha VOC B.1.1.7 strains (**Figure 1**). A 2 ml dose of 1×10^6^ TCID_50_ per animal was administered through intra-nasal (IN) and oral (PO) routes simultaneously. The remaining two non-infected sheep were separated and held in a dedicated clean room. At 1 day-post-challenge (DPC), the two naïve sheep were co-mingled with the principal infected animals as contact sentinels for the duration of the study. A subset of the principal infected sheep were euthanized and *postmortem* examination was performed at 4 (n=3) and 8 (n=3) DPC. *Postmortem* examination of the remaining two principal infected and two sentinel sheep was performed at 21 DPC (**Table 1**).

**Figure 1.**
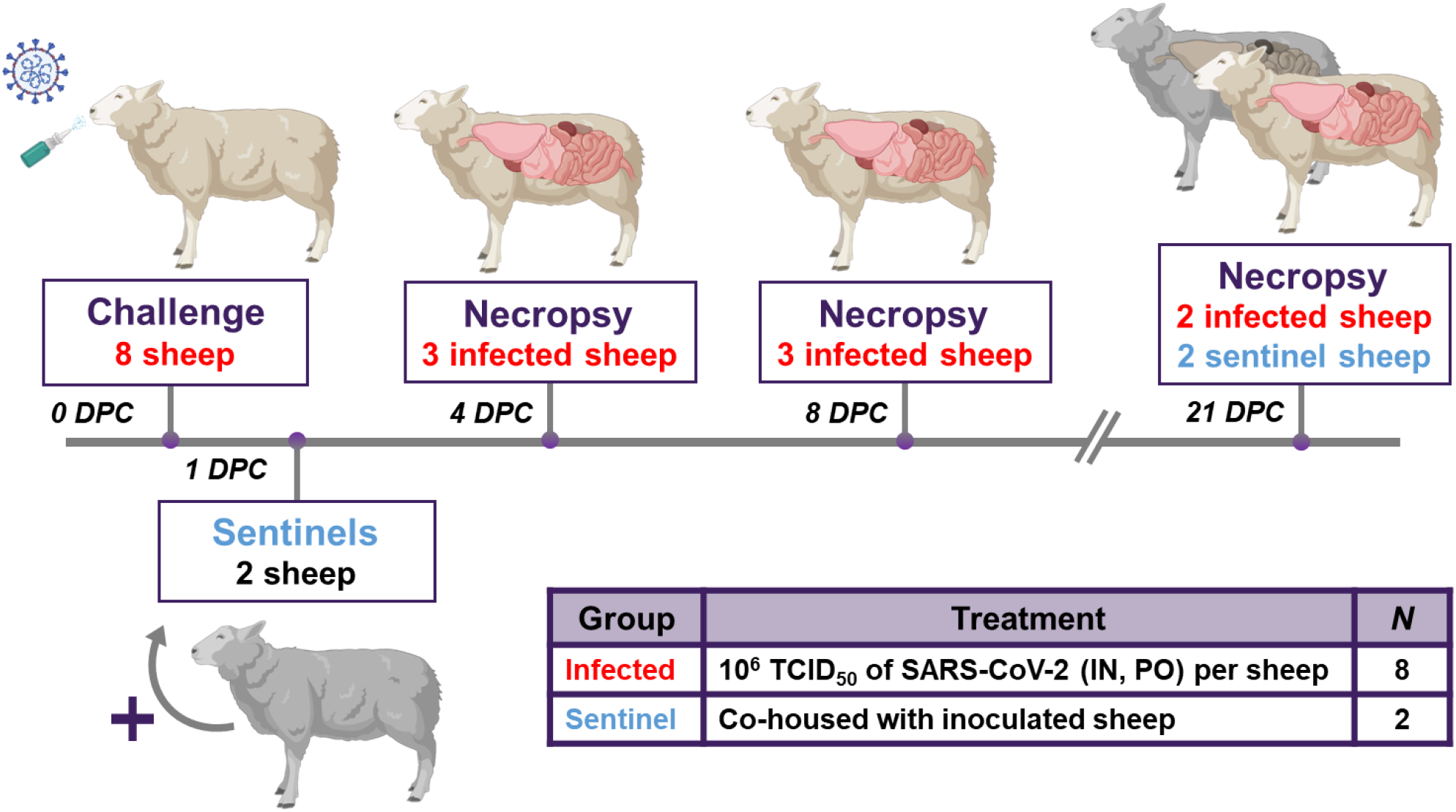
Study design. Eight sheep were inoculated with a mixture of SARS-CoV-2/human/USA/WA1/2020 lineage A (referred to as lineage A WA1; BEI item #: NR-52281) and SARS-CoV-2/human/USA/CA_CDC_5574/2020 lineage B.1.1.7 (alpha VOC B.1.1.7; NR-54011) acquired from BEI Resources (Manassas, VA, USA). A 2ml dose of 1×10^6^ TCID_50_ per animal was administered IN and PO. At 1-day post challenge (DPC), 2 sentinel sheep were co-mingled with the 8 principal infected animals to study virus transmission. Daily clinical observations and body temperatures were performed. Nasal/oropharyngeal/rectal swabs, blood/serum and feces were collected at 0, 1, 3, 5, 7, 10, 14 17 and 21 DPC. Postmortem examinations were performed at 4 (3 principal), 8 (3 principal) and 21 DPC (2 principal + 2 sentinels). BioRender.com was used to create figure illustrations.

**Table 1.**
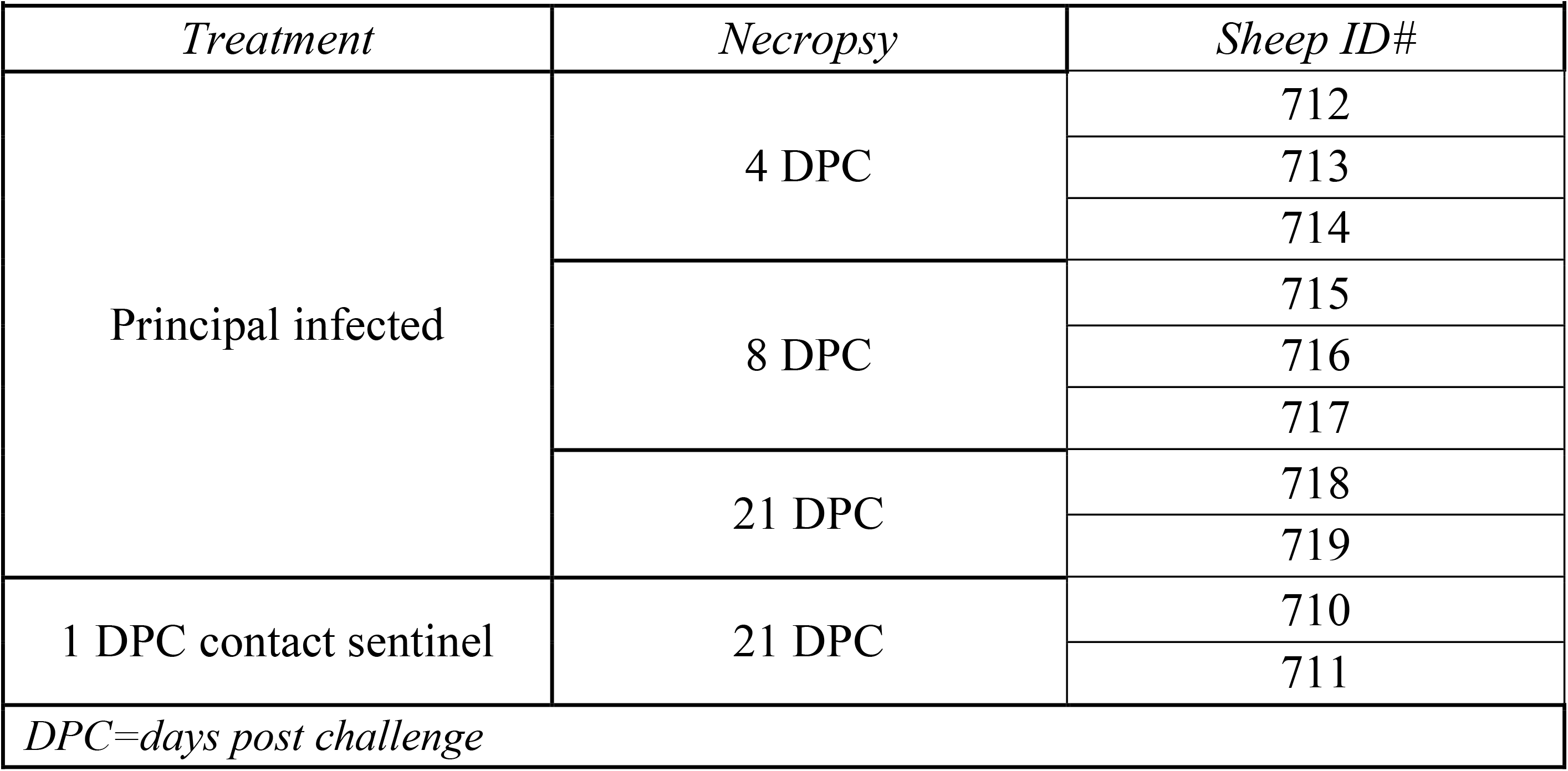
Animal treatment assignments.

### Clinical evaluations and sample collection

Sheep were observed daily for clinical signs. Clinical observations focused on activity level (response to human observer), neurological signs, respiratory rate, and presence of gastrointestinal distress. Rectal temperature, nasal, oropharyngeal, and rectal swabs were collected from animals at 0, 1, 3, 5, 7, 10, 14, 17 and 21 DPC. Swabs were placed in 2mL of viral transport medium (DMEM, Corning; combined with 1% antibiotic-antimycotic, ThermoFisher), vortexed, and aliquoted directly into cryovials and RNA stabilization/lysis Buffer RLT (Qiagen, Germantown, MD, USA). EDTA blood and serum were collected prior to challenge and on days 3, 7, 10, 14, 17, and 21 DPC. Full *postmortem* examinations were performed, and gross changes recorded. A comprehensive set of tissues were collected in either 10% neutral-buffered formalin (Fisher Scientific, Waltham, MA, USA), or as fresh tissues directly stored at −80°C. Tissues were collected from the upper respiratory tract (URT) and lower respiratory tract (LRT), central nervous system (brain and cerebral spinal fluid [CSF]), gastrointestinal tract (GIT) as well as accessory organs. The lungs were removed *in toto* including the trachea, and the main bronchi were collected at the level of the bifurcation and at the entry point into the lung lobe. Lung lobes were evaluated based on gross pathology and collected and sampled separately. Nasal wash and bronchoalveolar lavage fluid (BALF) were also collected during *postmortem* examination. Fresh frozen tissue homogenates were prepared as described previously (29). All clinical samples (swabs, nasal washes, BALF, CSF) and tissue homogenates were stored at −80°C until further analysis.

### RNA extraction and reverse transcription quantitative PCR (RT-qPCR)

SARS-CoV-2 specific RNA was detected and quantified using a quantitative reverse transcription real time – PCR (RT-qPCR) assay specific for the N2 segment as previously described (29). Briefly, nucleic acid extractions were performed by combining equal amounts of Lysis Buffer RLT (Qiagen, Germantown, MD, USA) with supernatant from clinical samples (swabs, nasal washes, BALF, CSF), tissue homogenates in DMEM (20% W/V), EDTA blood or body fluids. Sample lysates were vortexed and 200 μL was used for extraction using a magnetic bead-based extraction kit (GeneReach USA, Lexington, MA) and the Taco™ mini nucleic acid extraction system (GeneReach) as previously described (29). Extraction positive controls (IDT, IA, USA; 2019-nCoV_N_Positive Control), diluted 1:100 in RLT lysis buffer, and negative controls were included throughout this process.

Quantification of SARS-CoV-2 RNA was accomplished using an RT-qPCR protocol established by the CDC for detection of SARS-CoV-2 nucleoprotein (N)-specific RNA (https://www.fda.gov/media/134922/download). Our lab has validated this protocol using the N2 SARS-CoV-2 primer and probe sets (CDC assays for RT-PCR SARS-CoV-2 coronavirus detection, IDT, idtdna.com) in combination with the qScript XLT One-Step RT-qPCR Tough Mix (Quanta Biosciences, Beverly, MA, USA), as previously described (29). Quantification of RNA copy number (CN) was based on a reference standard curve method using a 5-point standard curve of quantitated plasmid DNA containing the N-gene segment (IDT, IA, USA; 2019-nCoV_N_Positive Control). The equation from this standard curve was used to extrapolate CN values from a 10-point standard curve of viral RNA extracted from a USA/CA_CDC_5574/2020 cell culture supernatant. This 10-point standard curve of viral RNA was used as reference for CN quantification from unknown samples to more accurately quantify Ct values at the extremity of quantification. Each sample was run in duplicate wells and all 96-well plates contained duplicate wells of quantitated PCR positive control (IDT, IA, USA; 2019-nCoV_N_Positive Control, diluted 1:100) and four non-template control wells. A positive Ct cut-off of 38 cycles was used when both wells were positive, as this represented a single copy number/µL and the LOD for this assay. Samples with one of two wells positive at or under CT of 38 were considered suspect-positive. Data are presented as the mean of the calculated N gene CN per mL of liquid sample or per mg of 20% tissue homogenate.

### Next-Generation Sequencing

RNA extracted from cell culture supernatant (virus stocks), clinical swab/tissue homogenates, and clinical samples were sequenced by next generation sequencing (NGS) using an Illumina NextSeq platform (Illumina Inc.) to determine the genetic composition (% lineage) of viral RNA in each sample. SARS-CoV-2 viral RNA was amplified using the ARTIC-V3 RT-PCR protocol [Josh Quick 2020. nCoV-2019 sequencing protocol v2 (GunIt). Protocols.io https://gx.doi.org/10.17504/protocols.io.bdp7i5rn]. Library preparation of amplified SARS-CoV-2 DNA for sequencing was performed using a Nextera XT library prep kit (Illumina Inc.) following the manufacturer’s protocol. The libraries were sequenced on the Illumina NextSeq using 150 bp paired end reads with a mid-output kit. Reads were demultiplexed and parsed into individual sample files that were imported into CLC Workbench version 7.5 (Qiagen) for analysis. Reads were trimmed to remove ambiguous nucleotides at the 5’ terminus and filtered to remove short and low-quality reads. The consensus sequences of viral stocks used for challenge material preparation were found to be homologous to the original strains obtained from BEI (GISAID accession numbers: EPI_ISL_404895 (WA-CDC-WA1/2020) and EPI_ISL_751801 (CA_CDC_5574/2020). To determine an accurate relative percentage of each SARS-CoV-2 lineage in each sample, BLAST databases were first generated from individual trimmed and filtered sample reads. Subsequently, two 40-nucleotide long sequences were generated for each strain at locations that include the following gene mutations: Spike (S) A570D, S D614G, S H1118H, S ΔH69V70, S N501Y, S P681H, S 982A, S T716I, S ΔY145, Membrane (M) V70, Nucleoprotein (N) D3L, N R203KG204R, N 235F, and non-structural NS3 T223I, NS8 Q27stop, NS8 R52I, NS8 Y73C, NSP3 A890D, NSP3 I1412T, NSP3 T183I, NSP6 ΔS106G107F108, NSP12 P323L, NSP13 A454V and NSP13 K460R. A word size of 40 was used for the BLAST analysis to exclude reads where the target mutation fell at the end of a read or reads that partially covered the target sequence. The two sequences for each of the locations listed above, corresponding to either lineage A (USA-WA1/2020) or alpha B.1.1.7 VOC (USA/CA_CDC_5574/2020), were subjected to BLAST mapping analysis against individual sample read databases to determine the relative amount of each strain in the samples. The relative amount of each strain present was determined by calculating the percent of reads that hit each of the twenty-three target mutations. The percentages for all mutations were averaged to determine the relative amount of each strain in the sample. Samples with incomplete or low coverage across the genome were excluded from analysis.

### Virus neutralizing antibodies

Virus neutralizing antibodies in sera were determined using microneutralization assay as previously described (29). Briefly, heat inactivated (56°C/30 min) serum samples were subjected to 2-fold serial dilutions starting at 1:20 and tested in duplicate. Then, 100 TCID_50_ of SARS-CoV-2 virus in 100 μL DMEM culture media was added 1:1 to 100 μL of the sera dilutions and incubated for 1 hour at 37°C. The mixture was subsequently cultured on Vero-E6/TMPRSS2 cells in 96-well plates. The neutralizing antibody titer was recorded as the highest serum dilution at which at least 50% of wells showed virus neutralization based on the absence of CPE observed under a microscope at 72 h post infection.

### Detection of antibodies by indirect ELISA

Indirect ELISAs were used to detect SARS-CoV-2 antibodies in sera with nucleocapsid (N) and the receptor-binding domain (RBD) recombinant viral proteins, both produced in-house (29). Briefly, wells were coated with 100 ng of the respective protein in 100 μL per well coating buffer (Carbonate–bicarbonate buffer; Sigma-Aldrich, St. Louis, MO, USA). Following an overnight incubation at 4°C, plates were washed three times with phosphate buffered saline (PBS-Tween 20 [pH=7.4]; Millipore Sigma), blocked with 200 μL per well casein blocking buffer (Sigma-Aldrich) and incubated for 1 hour at room temperature (RT). Plates were subsequently washed three times with PBS-Tween-20 (PBS-T). Serum samples were diluted 1:400 in casein blocking buffer, then 100 μL per well was added to ELISA plates and incubated for 1 hour at RT. Following three washes with PBS-T, 100 μL of HRP-labelled Rabbit Anti-Sheep IgG (H+L) secondary antibody (VWR, Batavia, IL, USA) diluted 1:1000 (100ng/mL) was added to each well and incubated for 1 hour at RT. Plates were then washed five times with PBS-T and 100 μL of TMB ELISA Substrate Solution (Abcam, Cambridge, MA, USA) was added to all wells of the plate and incubated for 5 minutes before the reaction was stopped. The OD of the ELISA plates were read at 450 nm on an ELx808 BioTek plate reader (BioTek, Winooski, VT, USA). The cut-off for a sample being called positive was determined as follows: Average OD of negative serum + 3X standard deviation. Everything above this cut-off was considered positive. Indirect ELISA was used to detect bovine coronavirus (BCoV) antibodies in sera with Spike (S) recombinant viral protein (LSBio, Seattle, WA, USA) using the methods described above.

### Histopathology

Tissue samples from the respiratory tract (nasal cavity [rostral, middle and deep turbinates following decalcification with Immunocal™ Decalcifier (StatLab, McKinney, TX, for 4-7 days at room temperature), trachea, and lungs as well as various other extrapulmonary tissues (liver, spleen, kidneys, heart, pancreas, gastrointestinal tract [stomach, small intestine including Peyer’s patches and colon], cerebrum [including olfactory bulb], tonsils and numerous lymph nodes) were routinely processed and embedded in paraffin. Four-micron tissue sections were stained with hematoxylin and eosin following standard procedures. Two independent veterinary pathologists (blinded to the treatment groups) examined the slides and morphological descriptions were provided.

### SARS-CoV-2-specific immunohistochemistry (IHC)

IHC was performed as previously described (29) on four-micron sections of formalin-fixed paraffin-embedded tissue mounted on positively charged Superfrost® Plus slides and subjected to IHC using a SARS-CoV-2-specific anti-nucleocapsid rabbit polyclonal antibody (3A, developed by our laboratory) with the method previously described (30). Lung sections from a SARS-CoV-2-infected hamster were used as positive assay controls.

## Results

### SARS-CoV-2 replication in ruminant-derived cell lines

Ruminant cell cultures derived from cattle (*Bos taurus*), sheep (*Ovis aries*), and pronghorn (*Antilocapra americana*) were tested for susceptibility to SARS-CoV-2 and viral growth kinetics (**Figure 2**). SARS-CoV-2 lineage A WA1 strain was found to replicate in both, primary and immortalized sheep kidney cell cultures with virus titers increasing over the course of 6 to 8 days post infection (DPI). SARS-CoV-2 did not replicate in the primary pronghorn lung cell cultures, or in the two immortalized bovine cell lines, a bovine kidney and a bovine fetal fibroblast cell; instead reduction of virus titer was observed by 2 DPI to 6 DPI (**Figure 2**). These results indicate that sheep may be susceptible to SARS-CoV-2 infection, while cattle and American pronghorn likely are not.

**Figure 2.**
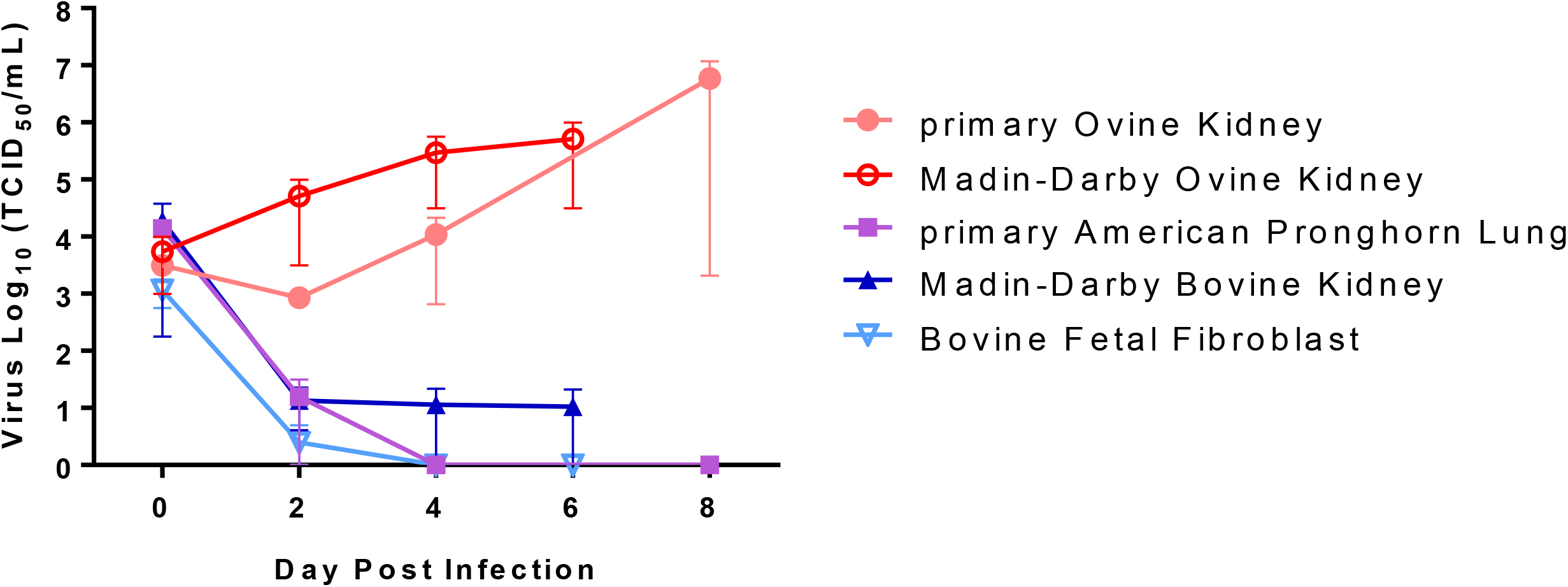
SARS-CoV-2 replication in ruminant-derived cell cultures. Primary ovine kidney cells, Madin-Darby ovine kidney, primary American pronghorn lung cells, bovine fetal fibroblast (BFF) and Madin-Darby bovine cell lines were infected with SARS-CoV-2 USA-WA1/2020 at 0.1 MOI and cell supernatants collected at 0, 2, 4, 6 or 8 days post infection (DPI). Cell supernatants were titrated on Vero E6 cells to determine virus titers. Mean titers and SEM of at least two independent infection experiments per cell line are shown.

### Sheep remain subclinical following challenge with SARS-CoV-2

Eight sheep were infected through IN and PO routes simultaneously with 1 ×10^6^ TCID_50_ per animal of a 1:10 titer ratio of the lineage A WA1 and the B.1.1.7-like alpha VOC strains. One day later, 2 sentinel sheep were co-mingled with the principal infected animals (**Figure 1**). Average daily rectal temperatures of all ten sheep showed that they did not become febrile after challenge or during the 21-day study (**Supplementary Figure 1**). In addition, no obvious clinical signs were observed in any of the principal infected or sentinel sheep. No weight loss, lethargy, diarrhea, inappetence, or respiratory distress was observed during the study.

### SARS-CoV-2 shedding from infected sheep

Nasal, oropharyngeal, and rectal swabs were collected over the course of the 21-day study and tested for the presence of SARS-CoV-2 RNA (**Figure 3**) and infectious virus (**Supplementary Table 1**). Viral RNA was detected in the nasal swab samples from 7 (ID#s 713-719) of the 8 principal infected sheep at 1 DPC, and only in one principal animal (#715) at 3 DPC. One oropharyngeal swab from a principal infected animal (#719) was also positive at 1 DPC. No other nasal or oropharyngeal swabs, and none of the rectal swab samples, collected up to 21 DPC were positive for viral RNA. Swab samples positive by RT-qPCR were also tested for the presence of infectious virus on susceptible VeroE6/TMPRSS2 cells, but no infectious virus was isolated. Nasal, oropharyngeal, and rectal swabs from the two sentinel sheep remained PCR negative over the course of 20 days of co-mingling with the principal infected animals.

**Figure 3.**
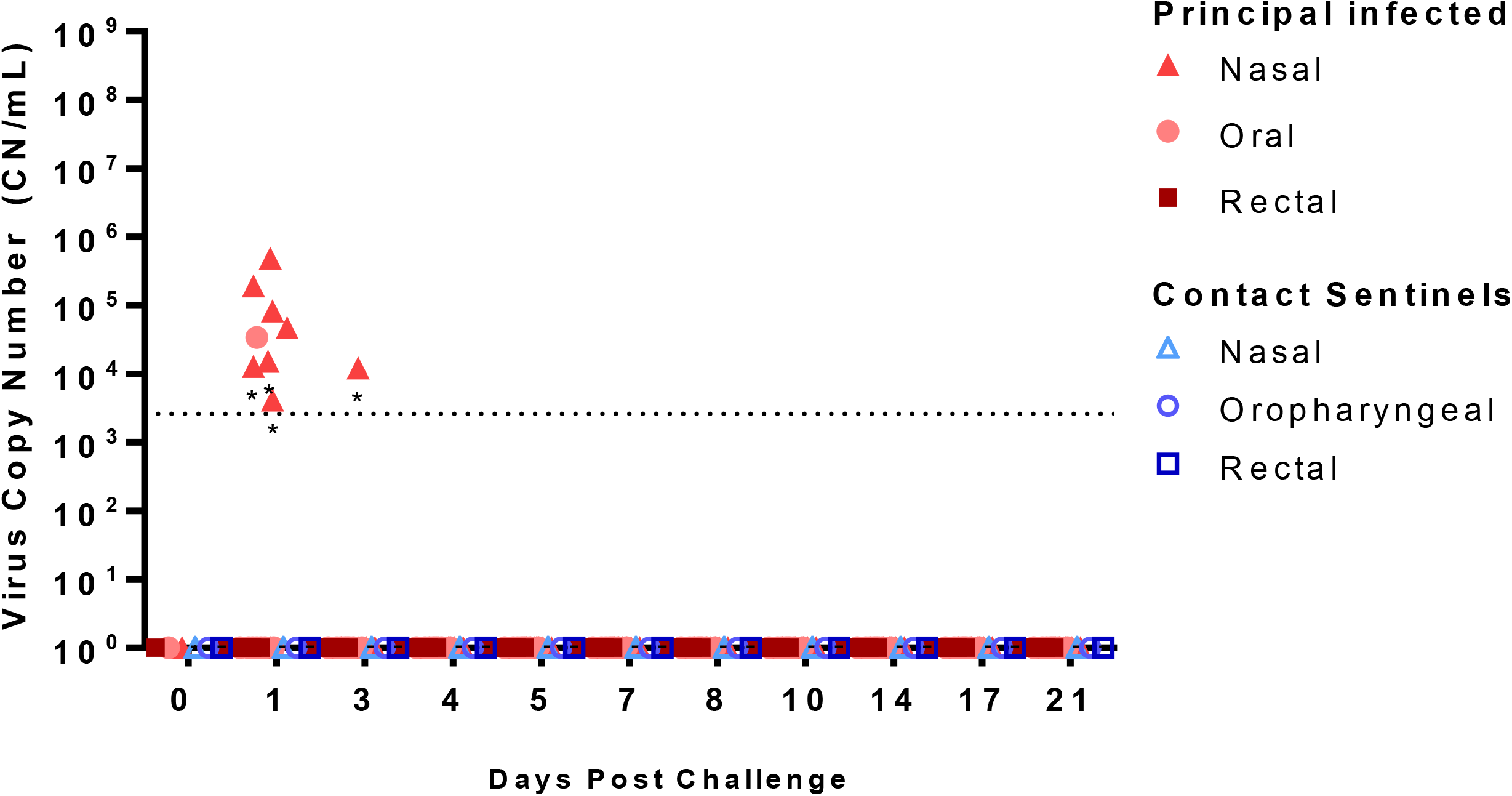
Viral RNA shedding of SARS-CoV-2-infected sheep. RT-qPCR was performed on nasal, oropharyngeal, and rectal swabs collected from principal infected (solid red symbols) and sentinel sheep (open blue symbols) on the indicated days post-challenge (DPC). Mean (n=2) viral RNA copy number (CN) per mL based on the SARS-CoV-2 nucleocapsid gene are plotted for individual animals. Asterisks (*) indicate samples with one out of two RT-qPCR reactions above the limit of detection, which is indicated by the dotted line.

### SARS-CoV-2 RNA detection in tissues of infected sheep

Tissues were collected from sheep euthanized at 4, 8, and 21 DPC (**Table 1**). SARS-CoV-2 RNA was detected in the respiratory tract tissues of all three principal infected sheep euthanized at 4 DPC, with the highest viral RNA loads detected in the trachea followed by the nasopharynx (**Figure 4A**). At 8 DPC, viral RNA was detected in some of the respiratory tissues of the three principal infected sheep, and similarly as found on 4 DPC, the nasopharynx and trachea had the highest RNA levels (**Figure 4B**). At 21 DPC, viral RNA was detected in the nasopharynx of both principal infected sheep euthanized at this time point, as well as in the conchae and ethmoturbinates of one principal infected animal (#718) (**Figure 4C**). On day 21 DPC, both sentinel sheep had viral RNA present in the conchae, and one sentinel animal (#710) also had detectable viral RNA in the trachea and bronchi (**Figure 4C**).

**Figure 4.**
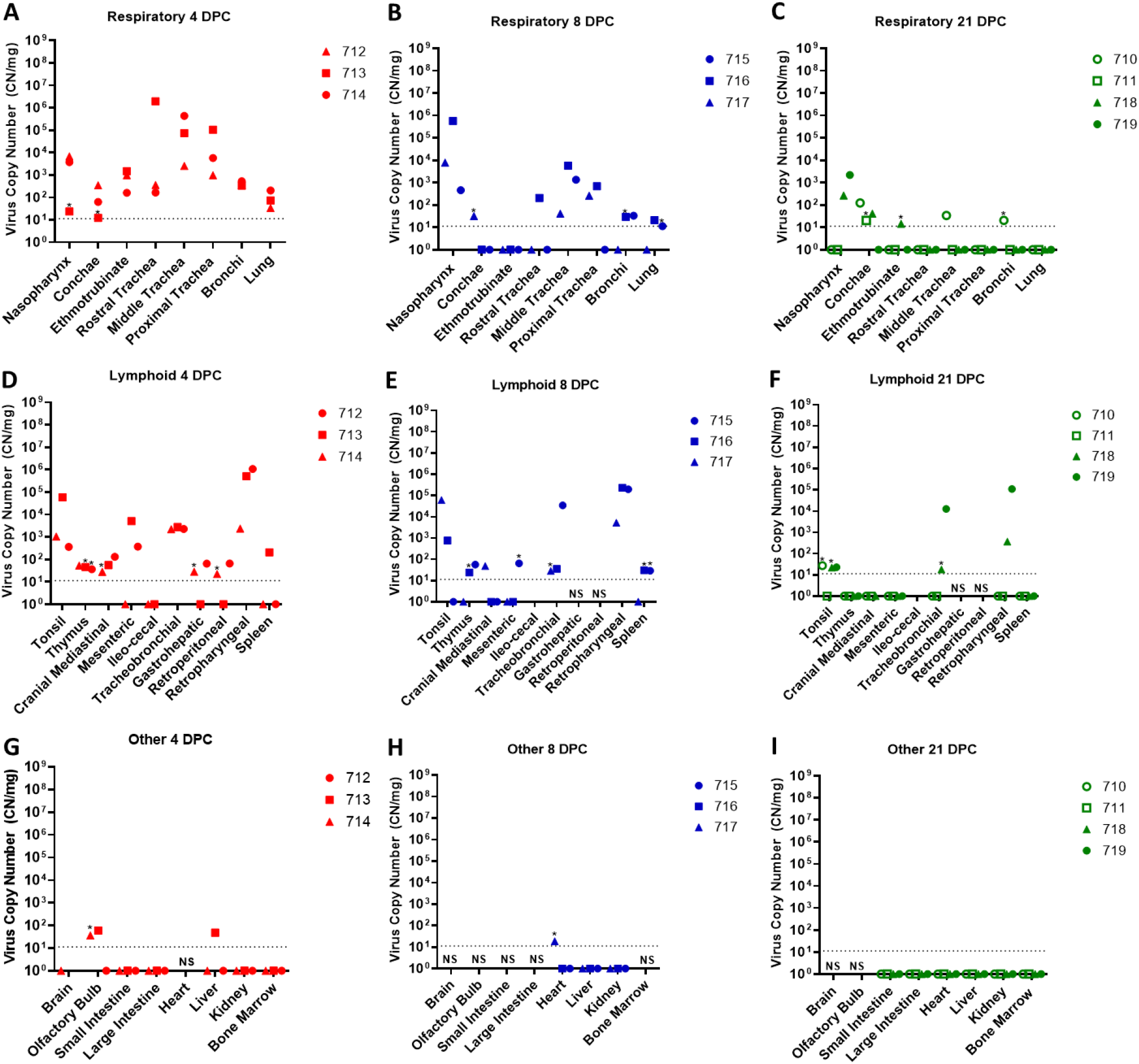
Viral RNA detected in tissues of SARS-CoV-2-infected sheep. RT-qPCR was performed on respiratory (A-C), lymphoid (D-F) and other (G-I) tissues of sheep euthanized at 4 (A,D,G), 8 (B,E,H), and 21 (C,F,I) days post challenge (DPC) to detect the presence of SARS-CoV-2-specific RNA. Mean (n=2) viral RNA copy number (CN) per mg of tissue based on the SARS-CoV-2 nucleocapsid gene are plotted for individual animals. Asterisks (*) indicate samples with one out of two RT-qPCR reactions above the limit of detection, which is indicated by the dotted line. NS = no sample. Solid symbols indicate principal animals necropsied at 4, 8, and 21DPC; open symbols indicate sentinel animals necropsied at 21 DPC.

SARS-CoV-2 RNA was also detected in the lymphoid tissues. At 4 DPC, the tonsil, thymus and several lymph nodes (cranial mediastinal, gastro-hepatic, retroperitoneal, retropharyngeal) from all three principal sheep euthanized on that day were RT-PCR positive (**Figure 4D**). The spleen, and the mesenteric, ileocecal and tracheobronchial lymph nodes of at least 1 or 2 out of the 3 principal sheep were also RNA positive at 4 DPC. At 8 DPC, viral RNA was detected in the lymphoid tissues of only some of the three principal sheep (**Figure 4E**). At 21 DPC, both principal infected sheep had detectable viral RNA in the tonsil and in the retropharyngeal and tracheobronchial lymph nodes; the tonsil of one sentinel (#710) was also RNA positive (**Figure 4F**). The lymphoid tissues with the highest viral RNA levels at 4, 8, and 21 DPC were the tonsil, and the retropharyngeal and tracheobronchial lymph nodes.

Other tissues such as the brain (only #714), olfactory bulb, small and large intestine, heart, liver, kidney and bone marrow were also collected at necropsy and tested for presence of SARS-CoV-2-specific RNA. At 4 DPC, the olfactory bulb of two of the three principal animals and the liver of animal #713 were RNA positive, whereas the small and large intestine, kidney, and bone marrow were negative (**Figure 4G**). At 8 DPC, the heart of sheep #717 was considered a suspect RNA positive, and the liver and kidney of all three principal sheep were negative (**Figure 4H**). At 21 DPC, the olfactory bulb, small and large intestine, heart, liver, kidney, and bone marrow from both principal and sentinel sheep were all negative for the presence of viral RNA (**Figure 4I**). In addition, whole blood was collected during the 21-day study, but no viral RNA was detected in the blood of any of the sheep enrolled in the study (data not shown).

Nasal washes, BALF, and CSF collected from sheep at necropsy were also tested for the presence of viral RNA. SARS-CoV-2 RNA was detected in the nasal washes of all principal infected sheep at 4 DPC (**Table 2**). Viral RNA was detected in the BALF of two out of the three principal sheep at 4 DPC (#713, 714) and in one animal at 8 DPC (#716). Nasal washes and BALF collected from principal and sentinel sheep at 21 DPC were all negative. No viral RNA was detected in the CSF of any of the sheep at any time point.

**Table 2.** Viral RNA detected in biological fluids collected from SARS-CoV-2 infected sheep.

|  |  |  |  |
| --- | --- | --- | --- |
| *Necropsy* | *Sheep ID* | *Biological* | *Mean CN/ML* |
| 4 DPC | 712 | BALF | ND |
|  |  | Nasal wash | 8.21E+04 |
|  |  | CSF | ND |
|  | 713 | BALF | 3.32E+03 |
|  |  | Nasal wash | 1.35E+05 |
|  |  | CSF | ND |
|  | 714 | BALF | 5.45E+04 |
|  |  | Nasal wash | 3.37E+04 |
|  |  | CSF | ND |
| 8 DPC | 715 | BALF | ND |
|  |  | Nasal wash | ND |
|  |  | CSF | ND |
|  | 716 | BALF | 3.08E+03* |
|  |  | Nasal wash | ND |
|  |  | CSF | ND |
|  | 717 | BALF | ND |
|  |  | Nasal wash | ND |
|  |  | CSF | ND |
| 21 DPC | 710 | BALF | ND |
|  |  | Nasal wash | ND |
|  |  | CSF | ND |
|  | 711 | BALF | ND |
|  |  | Nasal wash | ND |
|  |  | CSF | ND |
|  | 718 | BALF | ND |
|  |  | Nasal wash | ND |
|  |  | CSF | ND |
|  | 719 | BALF | ND |
|  |  | Nasal wash | ND |
|  |  | CSF | ND |

*\*1 out of 2 RT-qPCR reactions above the limit of detection; DPC=days post challenge; CN=RNA copy number; ND=not detected; BALF=bronchoalveolar lavage fluid; CSF=brain and cerebral spinal fluid*

Virus isolation was attempted on samples that were RT-qPCR positive having at least 10^3^ RNA copy number/mL (**Supplementary Table 1**). Viable virus was detected in proximal and rostral trachea samples (5×10^0^ TCID_50_/ml each) collected from principal infected sheep #713 at 4 DPC. No other samples had detectable viable virus.

### Serology

Indirect ELISA tests were used to detect SARS-CoV-2 antibodies against the recombinant N and the spike RBD antigens. Sera collected from principal infected sheep had detectable antibodies to N (**Figure 5A**) and RBD (**Figure 5B**) at 10, 14, 17 and 21 DPC. Sentinels did not develop antibodies to N or RBD above the assay cutoff. Only one of the principal infected sheep (#719) developed a low level of neutralizing antibodies with a 1:20 titer, detectable at 10 and 21 DPC (**Figure 5C**). Sera collected from sheep prior to SARS-CoV-2 challenge was also tested for reactivity against the bovine coronavirus spike antigen by indirect ELISA and were found to be negative (**Figure 5D**).

**Figure 5.**
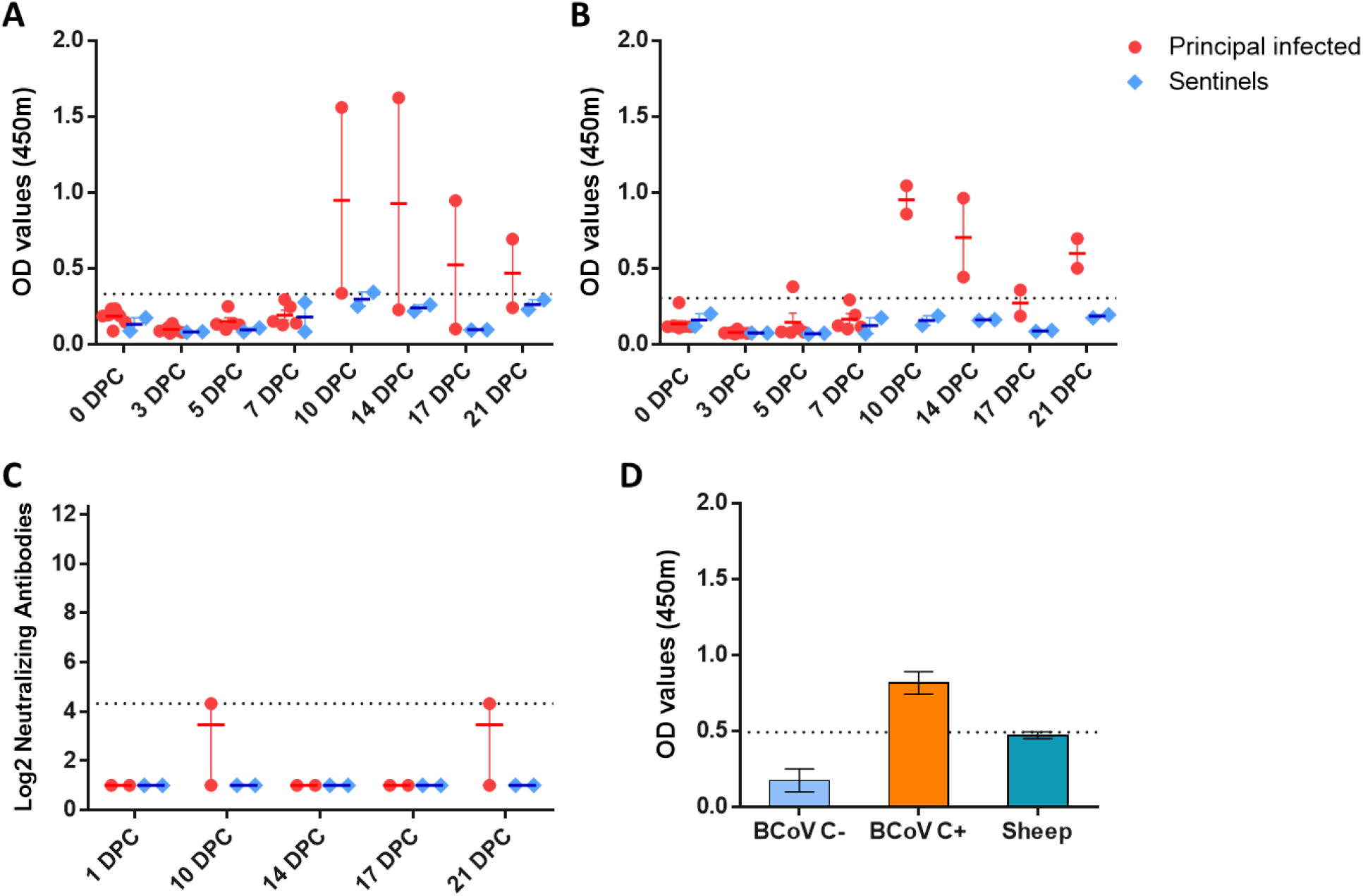
Serology of SARS-CoV-2 infected sheep. Detection of SARS-CoV-2 nucleocapsid protein (**A**), and the receptor binding domain (**B**) by indirect ELISA tests. The cut-off was determined by averaging the OD of negative serum + 3X the standard deviation as indicated by the dotted line. All samples with resulting OD values above this cut-off were considered positive. (**C**) Virus neutralizing antibodies detected in serum are shown as log2 of the reciprocal of the neutralization serum dilution. Sera were tested starting at a dilution of 1:20 which a cut-off of 1:10 indicated by the dotted line. (**D**) Sera from principal infected (n=8) and sentinel sheep (n=2) were tested against the bovine coronavirus (BCoV) spike protein using an indirect ELISA; both, positive (C+) and negative (C-) bovine control sera were included. The cut-off was determined by averaging the OD of negative serum + 3X the standard deviation as indicated by the dotted line. **A-D:** Mean with SEM are shown.

### Pathology

*Postmortem* pathological evaluations of sheep were performed at 4, 8, and 21 DPC (**Table 1**). Overall, no significant gross lesions were observed. Prominent respiratory tract-associated lymphoid tissue was noted in the upper and lower respiratory tract of all infected sheep at 4 and 8 DPC as well as mock-infected controls (**Figures 6 and 7**). These lymphoid aggregates were subjectively more frequent and prominent at 8 DPC compared to 4 DPC. No other changes were noted in nasal turbinates or lungs at 4 DPC, and no viral antigen was detected in these organs (**Figure 6 A, B, E and F**). In addition to the prominent lymphoid aggregates, moderate tracheitis was observed in animals #713 and #714 at 4 DPC, with evidence of lymphocytic transmigration and few scattered necrotic epithelial cells (**Figure 6C**). Viral antigen was detected in the trachea of sheep #713, and was solely localized to lymphoid aggregates in the lamina propria and, based on the morphology of the immunopositive cells, these likely represent macrophages/dendritic cells (i.e., antigen presenting cells) (**Figure 6D**). Lymph nodes associated with the respiratory tract, tonsils, and third eyelids were characterized by severe lymphoid hyperplasia with traces of viral antigen detected in few cells resembling macrophages/dendritic cells, within a few lymphoid aggregates of the pharyngeal region of animals #712 and #713 euthanized on 4 DPC. Overall, viral antigen appeared associated with lymphoid aggregates, with limited antigen positivity observed primarily in phagocytic cells. No viral antigen was detected within the respiratory epithelium, associated glands, or pulmonary pneumocytes.

**Figure 6.**
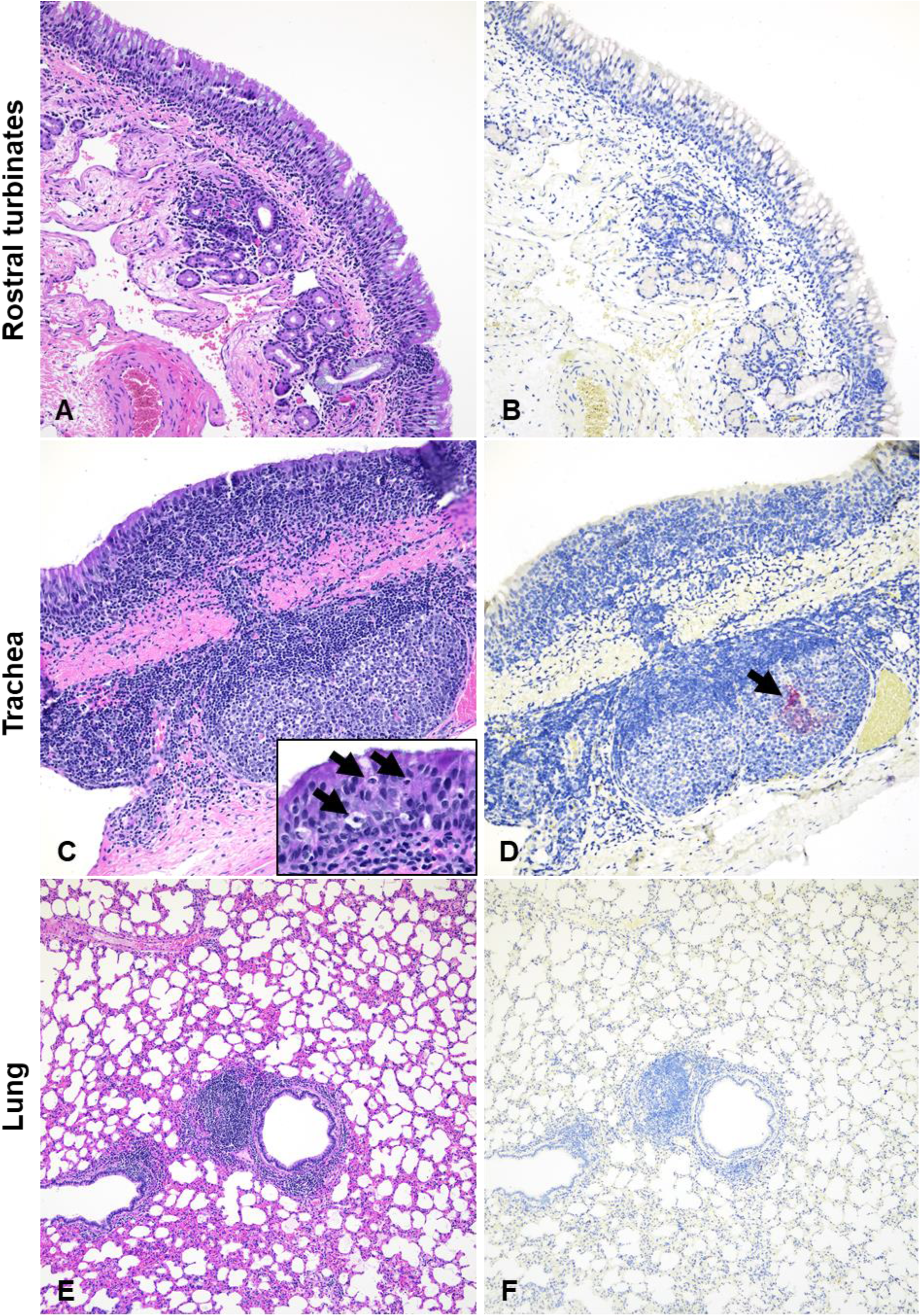
Histopathology and SARS-CoV-2 antigen distribution in the upper and lower respiratory tract of infected sheep at 4 DPC. Rostral turbinates (A and B), trachea (C and D), and lung (E and F). At 4 DPC, minimal changes were noted in the rostral turbinates, with mild, dispersed and aggregates of lymphocytes. No viral antigen was detected (B). In the trachea, there were multifocal prominent aggregates of lymphocytes and plasma cells in the lamina propria and extending/transmigrating through the lining epithelium, with few individual cell degeneration and necrosis (C and inset [arrows]). Sporadic lymphoid aggregates showed viral antigen (D, arrow). In the pulmonary parenchyma, bronchioles and blood vessels were delimited by hyperplastic bronchus-associated lymphoid tissue (BALT) (E), no viral antigen was detected (F). H&E and Fast Red, 200X total magnification (A-D) and 100X total magnification (E and F).

No significant histologic changes were noted in the respiratory tract at 8 DPC other than the hyperplastic respiratory tract-associated lymphoid tissue (**Figure 7**). In a single animal (#716), there was moderate lymphocytic tracheitis with minimal adenitis, transmigration of lymphocytes along the lining epithelium, and occasionally necrotic epithelial cells. No viral antigen was detected in the respiratory tract (nasal passages, trachea, and lungs) or associated lymphoid tissue at 8 DPC.

**Figure 7.**
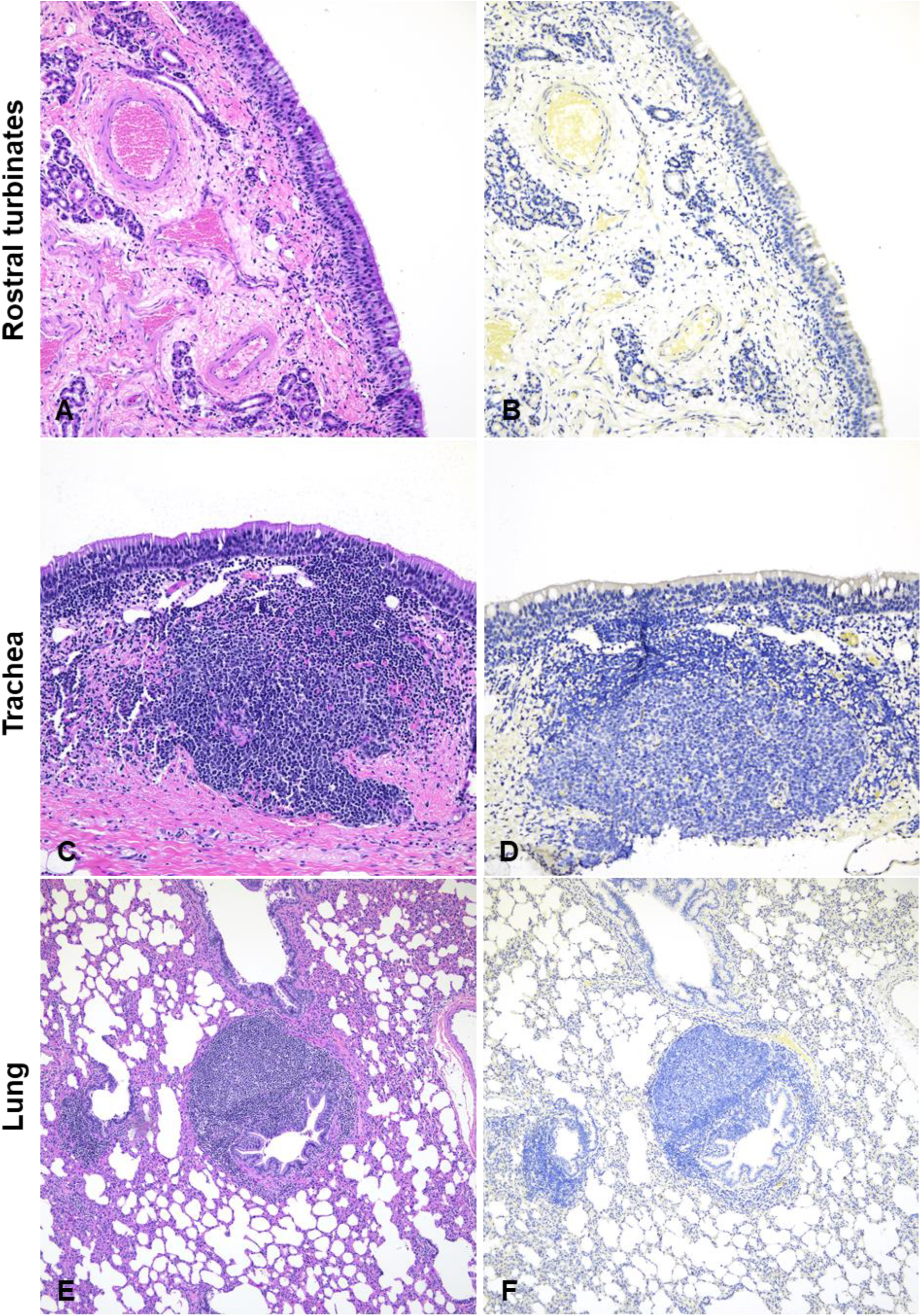
Histopathology and SARS-CoV-2 antigen distribution in the upper and lower respiratory tract of infected sheep at 8 DPC. Rostral turbinates (A and B), trachea (C and D), and lung (E and F). At 8 DPC, rostral turbinates were within normal limits and no viral antigen was detected (A and B). The tracheal lamina propria had multiple prominent and dense lymphoid aggregates (C) but no evidence of viral antigen (D) or epithelial alterations. In the pulmonary parenchyma, bronchioles and blood vessels were frequently delimited by prominent BALT) (E), no viral antigen was detected (F) H&E and Fast Red, 200X total magnification (A-D) and 100X total magnification (E and F).

### SARS-CoV-2 competition in co-infected sheep

To study the competition of two SARS-CoV-2 strains, the sheep challenge inoculum was prepared as a mixture of two SARS-CoV-2 isolates which were representative of the ancestral lineage A and the B.1.1.7-like alpha VOC. Next generation sequencing (NGS) was used to determine the percent presence of each strain in various swab and tissue samples collected from each sheep. The intention was to use a 1:1 titer ratio of each strain to inoculate sheep; however, back-titration showed that the actual ratio was closer to 1:10 of the WA1 lineage A to the B.1.1.7-like alpha VOC. The overall results indicate that the SARS-CoV-2 alpha variant outcompeted the ancestral lineage A strain in sheep (**Table 3**). NGS analysis consisted of 1 DPC nasal swabs, and various respiratory and lymphoid tissues collected at 4, 8 and 21 DPC. Analysis of the 1 DPC nasal swab samples of principal infected sheep #715 and #719 showed 100% presence of the B.1.7-like alpha VOC. The alpha VOC was found at 99.7-100%, compared to 0.0-0.3% of the ancestral lineage A strain in respiratory tissues (ethmoturbinates, nasopharynx, and trachea) of principal infected sheep #712, 713, and 714 analyzed at 4 DPC. At 8 DPC, the alpha VOC was found at 99.1-100% compared to 0.0-0.9% of the ancestral lineage A strain in the nasopharynx and trachea of principal infected sheep #715, 716, and 718. In the tonsil, the B.1.1.7-like alpha VOC was present at 76.4-100% in all three of the principal infected sheep euthanized at 4 DPC, and at 98.6-99.9% in 2 out of 3principal infected sheep at 8 DPC. This indicates that there was some replication of the lineage A strain in at least some of the tissues, especially the tonsil, of some of the sheep. No sequencing data is available for samples collected from principal infected or sentinel sheep at 21 DPC, due to low RNA copy numbers in the tissues or mapped reads.

**Table 3.**
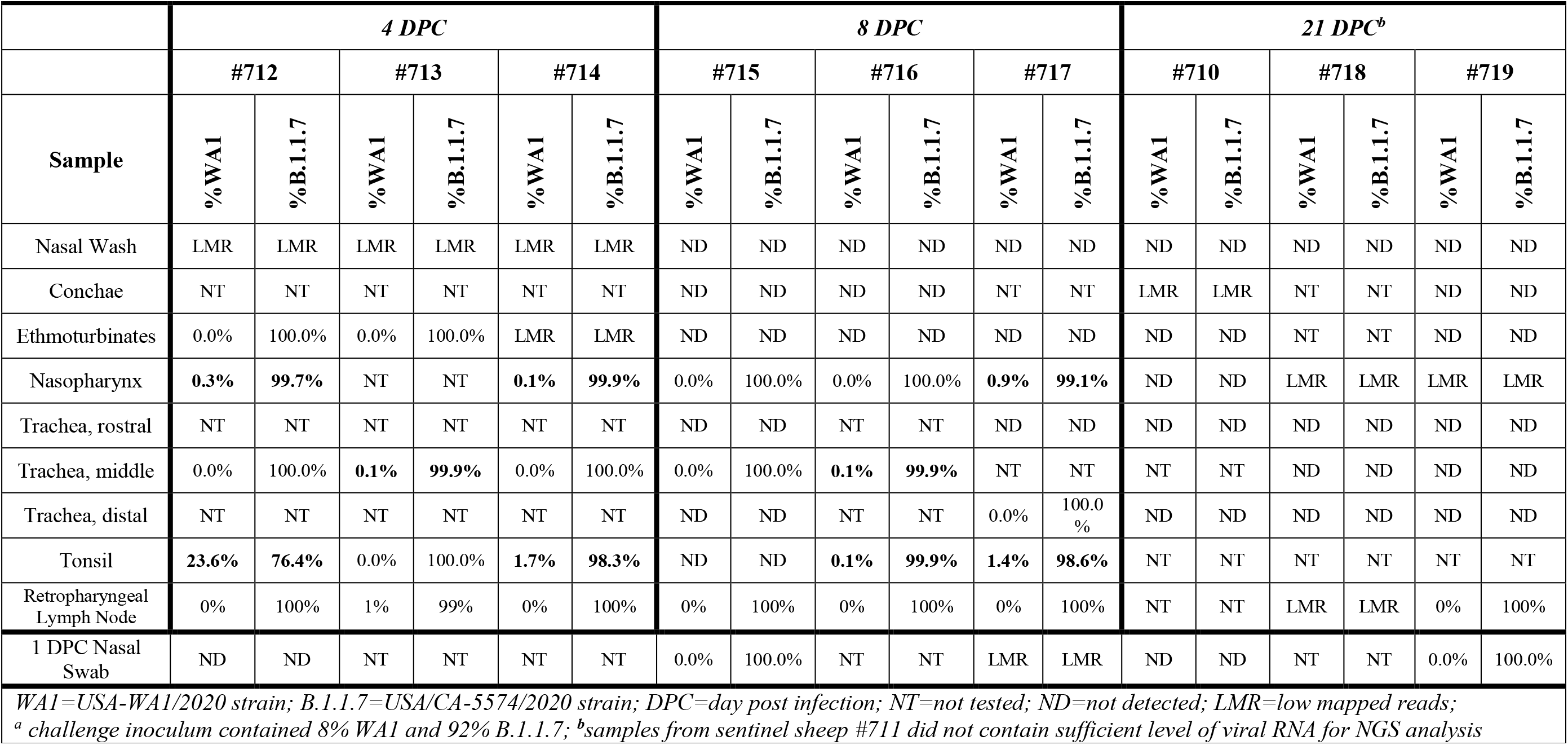
SARS-CoV-2 competition in sheep co-infected^a^ with lineage A and B.1.1.7 strains determined by next generation sequencing.

## Discussion

The emergence, rapid evolution, and persistence of SARS-CoV-2 has been an unwelcome reminder of our co-existence with, and vulnerability to, formidable microorganisms. It has been shown that emerging infectious diseases (EID) of humans frequently arise from the human-animal interface, i.e. are zoonotic pathogens (31, 32). Some EID’s resolve themselves in dead-end hosts while other pathogens, such as SARS-CoV-2, adapt quickly to new hosts and maintain transmission cycles in susceptible populations. The public health implications of SARS-CoV-2 do not start or stop with humans. Similar to SARS-CoV, which emerged in 2002, SARS-CoV-2 utilizes the ACE2 receptor to bind to and enter host cells. Provided the high conservation of ACE2 receptors amongst mammalian species, many animal species are potentially susceptible to SARS-CoV-2 infection (24). Therefore, a holistic, *One-Health* approach is necessary to fully understand and properly mitigate further escalation of the SARS-CoV-2 pandemic and its impacts.

So far, the susceptibility and epidemiological role of domestic ruminant species has been largely understudied. Sheep are a valuable agricultural species and are in close contact with potential SARS-CoV-2 reservoir species including humans, cats, deer, mustelids and rodents. Current data regarding their susceptibility is inconclusive. In order to address this gap, we investigated the susceptibility of sheep to SARS-CoV-2 by *in vitro* infection of sheep- and other ruminant-derived cell cultures and by *in vivo* challenge of sheep with SARS-CoV-2. In addition, we introduced two sentinel sheep to evaluate the potential of transmission of the virus from principal infected animals to naïve sheep. Furthermore, we co-infected sheep with the ancestral lineage A and the B.1.1.7-like alpha VOC SARS-CoV-2 strains in order to study virus strain competition in the animal host.

Our results of SARS-CoV-2 infected ruminant-derived cell cultures showed that sheep primary kidney cells and an immortal sheep kidney cell line supported SARS-CoV-2 infection and replication, while the pronghorn and bovine cell cultures used in this study did not (**Figure 2**). These results are consistent with several previous *in silico, in vitro* and *in vivo* studies (17, 24, 25, 33). However, as we have not monitored levels of ACE2 on the cell surface of these cell lines, we cannot exclude that lack of SARS-CoV-2 infection in pronghorn and bovine cell cultures is due to lack of expression of the host ACE2 receptor in these cell lines. In fact, computational modeling studies by Damas et al. (24) predicted the ACE2 molecule of cattle (*Bos taurus*), sheep (*Ovis aries*) and pronghorn (*Antilocapra americana*) to have a medium binding score with the RBD region of the SARS-CoV-2 spike protein, identifying these ruminant species as potential susceptible hosts. Furthermore, results from a study with SARS-CoV-2 infection of tracheal and lung organ cultures from sheep and cattle *ex vivo* showed that both were capable of supporting viral replication (25). Together these studies and ours suggest that sheep seem to have low susceptibility toSARS-CoV-2 infection. However, a serological survey in Spain of sheep in frequent contact with humans during the pandemic did not provide evidence to support infection of sheep with SARS-CoV-2 (26).

Our results showing that the Madin-Darby bovine kidney (MDBK) cells and bovine fetal fibroblasts do not support SARS-CoV-2 infection are consistent with a study by Hoffman and colleagues (33) that utilized a Spike-based pseudovirus system that demonstrated MDBKs do not support SARS-CoV-2 cell entry. Furthermore, a study in cattle showed that only 2 out of 6 experimentally challenged animals became infected and that transmission of the virus did not occur to co-housed naïve animals (17). Collectively these studies indicate that susceptibility of cattle to SARS-CoV-2 infection is low.

In this study, we determined susceptibility and transmission in experimentally challenged sheep. Sheep remained subclinical throughout the 21-day long study, and the duration of SARS-CoV-2 RNA shedding in clinical samples was rather short and primarily from the nasal cavity. At 1 DPC viral RNA was detected in nasal swabs of seven out of eight principal infected sheep. It cannot be ruled out that this was artifact or residual from the challenge inoculum. However, one principal infected sheep (#715) had a RT-qPCR positive nasal swab at 3 DPC and virus was isolated and viral antigen detected in the trachea of sheep #713 at 4 DPC. These findings are indicative of limited virus replication. Viral RNA detected in the oral swabs was limited to only 1 animal at 1 DPC, and no viral RNA was detected from rectal swabs from any sheep during the 21-day study.

Viral RNA was frequently detected in the respiratory and lymphoid tissues at 4 and 8 DPC, and less frequently at 21 DPC. Low levels of viral RNA were also detected in the olfactory bulb, liver and heart in a few animals up to 8 DPC. Nasopharynx, trachea, tonsil, and tracheobronchial and retropharyngeal lymph node tissues had the highest viral RNA levels. In addition, viable virus was isolated from the trachea of one of the challenged animals at 4 DPC, evident of active SARS-CoV-2 infection. The prominent lymphoid hyperplasia along the respiratory tract was a feature common to both infected and mock-infected animals. Even though this response seems to be most prominent at 8 DPC, its background presence in mock-infected animals precludes establishment of a direct association with SARS-CoV-2 infection. Few animals at 4 DPC showed mild to moderate tracheitis with evidence of epithelial alterations, however, viral antigen was not detected in the respiratory epithelium and solely localized to phagocytic cells within lymphoid aggregates (#713). Overall, viral antigen appeared associated with lymphoid aggregates, with limited antigen positivity observed primarily in phagocytic cells. Alternatively, SARS-CoV-2 antigen might have been acquired by phagocytosis of viral proteins rather than by limited infection. This observation explains the lack of significant virus shedding and effective transmission to co-mingling animals. Together our results indicate that sheep can be experimentally infected with SARS-CoV-2 resulting in a limited infection primarily associated with the upper respiratory tract and regional lymphoid tissues. Domestic sheep showed low susceptibility to SARS-CoV-2 infection and limited ability to transmit to contact animals.

The two contact sentinel sheep did not shed viral RNA nor seroconverted during the 21-day long study, but low levels of viral RNA were detected at 21 DPC in the trachea, bronchi and tonsil of one sentinel sheep, and the conchae of both sentinel animals. This suggests that transmission could occur but was not very effective in order to result in detectable virus shedding or a robust immune response in the contact sentinel sheep, i.e. a productive SARS-CoV-2 infection was not established in the sentinel animals. While viral shedding from principal infected animals did not appear to be sufficient to cause a productive infection in naïve contact sheep in our study, transmission to other highly susceptible species such as humans or other animals could be of potential concern.

Finally, the virus competition results from co-challenge of sheep demonstrated co-infection by both SARS-CoV-2 strains and confirmed the competitive advantage of the SARS-CoV-2 B.1.1.7-like alpha VOC over the ancestral lineage A strain in sheep. However, the input ratio of the two virus strains was unintentionally biased toward the alpha VOC (10x), therefore limited conclusions can be drawn from this particular experiment. Nonetheless, these results confirm the ability of the SARS-CoV-2 alpha VOC to infect sheep and its increased replicative capacity in general.

In conclusion, our results demonstrate that experimental challenge of sheep with SARS-CoV-2 results in a limited subclinical infection, and while transmission to naïve co-mingled sheep appeared to occur, it did not lead to highly productive infection; therefore, domestic sheep are unlikely to be amplifying hosts for SARS-CoV-2. Based on our results and the currently available published data, further investigations into SARS-CoV-2 infection in sheep and other ruminant species are warranted. The identification of additional susceptible hosts provides critical information for SARS-CoV-2 epidemiology, in order to establish surveillance protocols, and to improve our mitigation strategies and preventative measures at the human-animal interface.

## Acknowledgments

We thank the staff of KSU Biosecurity Research Institute, the histology laboratory at the Kansas State Veterinary Diagnostic Laboratory (KSVDL), members of the Histology and Immunohistochemistry sections at the Louisiana Animal Disease Diagnostic Laboratory (LADDL), the Comparative Medicine Group staff at Kansas State University and technical support from Emily Gilbert-Esparza, Yonghai Li of KSU, and Jeana Owens and Dane Jasperson from USDA ARS. The SARS-CoV-2 strains USA/CA-5574/2020 and USA/WA1/2020 were obtained through BEI Resources (catalog # NR-52281 and #54011). We also thank Dr. Kyeong-Ok Chang for the Vero E6/TMPRSS2 cells used in these studies.

