## Supplemental Figures & Tables for "Susceptibility of sheep to experimental co-infection with the ancestral lineage of SARS-CoV-2 and its alpha variant" (from another version of this preprint: version 1, 2021-11-17)

**SUPPLEMENTARY FIGURES & TABLES**

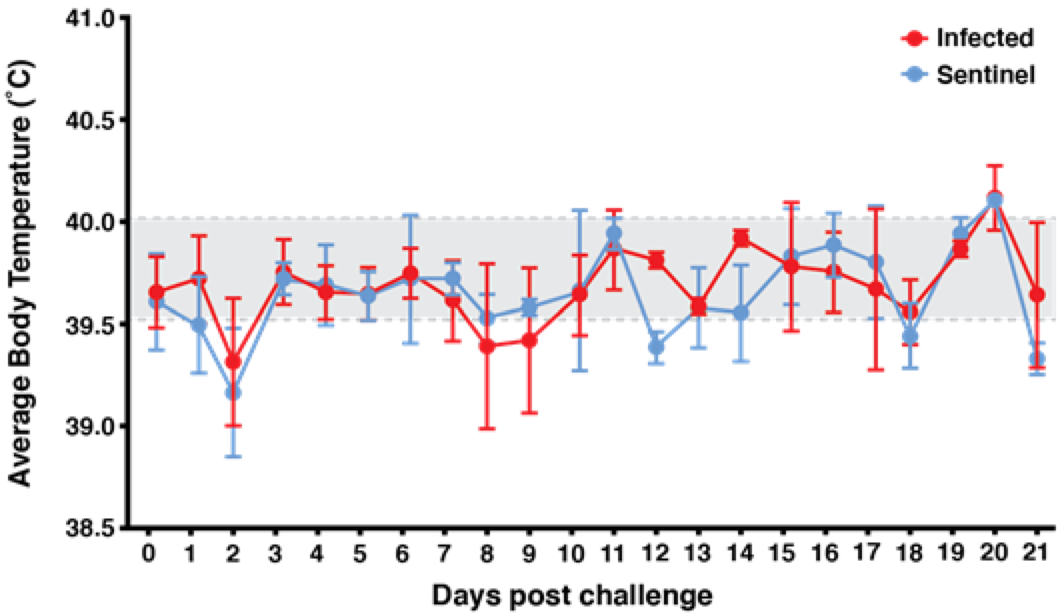


**Supplementary Figure 1. Body temperature.** Daily average rectal temperatures of sheep inoculated with SARS-CoV-2 (red) and co-mingled sentinel sheep (blue) showed no significant change over the course of the experiment. The baseline temperature (grey area; 39.5°C to 40°C) was determined from all sheep before infection.

| **Supplementary Table 1. Virus isolation from tissues tested.** | | | |
|---|---|---|---|
| *DPC* | *Sample Type* | *Animal Number* | *Virus titer ($TCID_{50}$/mL)* |
| 1 | Nasal swab | 713, 714, 715, 717, 719 | NEG |
| | Oral swab | 719 | NEG |
| 3 | Nasal swab | 715 | NEG |
| 4 | Nasal wash | 712, 713, 714 | NEG |
| | BALF | 714 | NEG |
| | Conchae | 712 | NEG |
| | Ethmoturbinates | 712, 713 | NEG |
| | Nasopharynx | 712, 714 | NEG |
| | Trachea | 712, 713, 714 | NEG, 5.00E+00*, NEG |
| | Bronchi | 712, 713, 714 | NEG |
| | Lung | 714 | NEG |
| | Tonsil | 712, 713, 714 | NEG |
| | Retropharyngeal lymph node | 712, 713 | NEG |
| | Tracheobronchial lymph node | 712, 713, 714 | NEG |
| | Mesenteric lymph node | 712, 713 | NEG |
| 8 | Nasopharynx | 715, 716, 717 | NEG |
| | Trachea | 715, 716, 717 | NEG |
| | Tonsil | 716, 717 | NEG |
| | Retropharyngeal lymph node | 715, 716, 717 | NEG |
| | Tracheobronchial lymph node | 715 | NEG |
| 21 | Nasopharynx | 719 | NEG |
| | Retropharyngeal lymph node | 718, 719 | NEG |
| | Tracheobronchial lymph node | 719 | NEG |
| **virus isolated from rostral and proximal trachea samples of sheep #713* | | | |
